# Dynamical Patterning Modules and Network Motifs as joint determinants of development: lessons from an aggregative bacterium

**DOI:** 10.1101/655381

**Authors:** Alejandra Guzmán-Herrera, Juan A. Arias Del Angel, Natsuko Rivera-Yoshida, Mariana Benítez, Alessio Franci

## Abstract

Development and evolution are dynamical processes under the continuous control of organismic and environmental factors. Generic physical processes, associated with biological materials and certain genes or molecules, provide a morphological template for the evolution and development of organism forms. Generic dynamical behaviors, associated with recurring network motifs, provide a temporal template for the regulation and coordination of biological processes. The role of generic physical processes and their associated molecules in development is the topic of the Dynamical Patterning Module (DPM) framework. The role of generic dynamical behaviors in biological regulation is studied via the identification of the associated Network Motifs (NM). We propose a joint DPM-NM perspective on the emergence and regulation of multicellularity focusing on a multicellular aggregative bacterium, *Myxococcus xanthus*. Understanding *M. xanthus* development as a dynamical process embedded in a physical substrate provides novel insights into the interaction between developmental regulatory networks and generic physical processes in the evolutionary transition to multicellularity.

## 1. Introduction

The transition from unicellular to multicellular life forms is one of the major evolutionary events, independently occurring multiple times and among different lineages (Maynard-Smith & Szathmáry, 1995; Grosberg & Strathmann, 2007; Bonner, 2016). The repeated occurrence of multicellularity enables the comparative and integrative study of potentially general molecular components, principles and mechanisms in development (Bonner, 2000). Such comparative studies may involve whole genomes, specific gene sequences, expression patterns and morphogenetic processes (*e.g.* Nanjundiah *et al.*, 2018; del Campo *et al.*, 2014; Zhu *et al.*, 2010). However, most of these studies have focused on organisms that develop multicellular structures by incomplete cell division (cells “staying together”, mainly in plants and animals), whereas aggregative organisms that exhibit another ubiquitous trajectory to multicellularity (cells “coming together”, as in some groups of fungi, bacteria and cellular slime molds) have been less studied (Figure 1).

**Figure 1.**
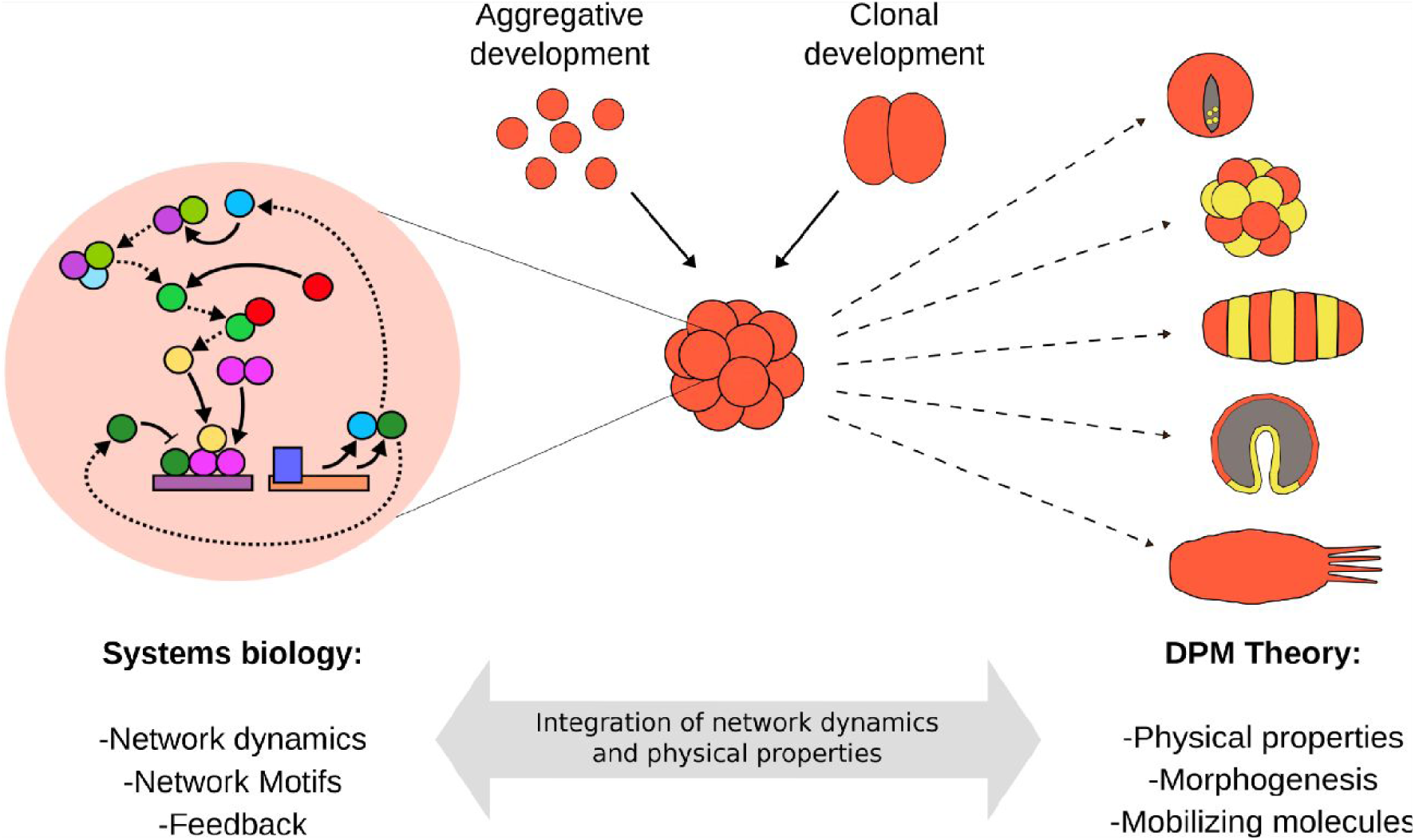
Comparison of DPM theory and Systems Biology. Independently of its origin (clonal or aggregative), multicellularity has emerged many times in the history of evolution. Two theories shed light on this Evo-Devo phenomenon: DPM theory (right) and Systems Biology (left). The two theories can be integrated to shed a more complete picture of the transition to multicellularity across-scales.

Studying the collective behaviors and patterning processes that occur in bacteria might be particularly illuminating since they take place at the spatial scale at which evolutionary transitions to multicellularity may have occurred. Such behaviors and patterns might be particularly evident in bacterial colonies, understood as discrete clusters of conspecific cells, that develop into multicellular structures with differentiated cells and stereotypic spatial arrangements (Bonner, 2000; Rivera-Yoshida *et al.*, 2018). *Myxococcus xanthus* is a delta-proteobacterium that enables the study of colonial developmental dynamics. This bacterium develops by aggregation into three-dimensional multicellular structures, called fruiting bodies, when nutrient availability in the medium is low. During its development, at least three different cell fates are observed: peripheral cells, stress-resistant spores and cells undergoing programmed cell death (Higgs *et al.*, 2014). Due to its reproducible developmental process, *M. xanthus* has been considered as a good model organism to investigate multicellular development and the evolution of multicellularity (Yang & Higgs *et al.*, 2014; Arias Del Angel *et al.*, 2017).

Both the regulatory network and the molecular mechanisms involved in *M. xanthus* development have been extensively studied (Kroos, 2007; Escalante *et al.*, 2012; Higgs *et al.*, 2014; Bretl & Kirby, 2016). The analysis of the structural and dynamical properties of this network has contributed to the understanding of the mechanisms underlying cell-fate determination in this bacterium (Arias Del Angel *et al.*, 2018, 2019). Furthermore, some studies have shown that *M. xanthus* development can be partially explained by physical and chemical morphogenetic processes affecting living and non-living matter, such as surface-tension-driven coarsening and substrate-mediated interactions (Bahar *et al.*, 2014; Rivera-Yoshida *et al.*, 2019). This plethora of factors and processes involved in *M. xanthus* aggregative development calls for integrative approaches that consider molecular network dynamics in association with morphogenetic processes.

In this study, we make use of two important complementary frameworks, Dynamical Patterning Modules (DPMs) (Newman & Bhat, 2008, 2009) and Network Motifs (NMs) (Alon, 2007), to analyze *M. xanthus* developmental molecular regulatory network (MRN). We briefly review the main strengths and weaknesses of each framework (section 2) before combining them into a DPM-NM approach that integrates the physical aspects of morphogenetic processes, mediated by DPMs, with their correct regulation and temporal coordination, mediated by NM dynamics. We use this integrated approach to dissect and analyze the dynamics of *M. xanthus* developmental MRN (section 3), highlighting the tight synergy between the two frameworks. Finally, we discuss the key points of our analysis, its importance for understanding multicellularity, and its relevance for evolutionary-developmental (Evo-Devo) biology (section 4).

## 2. DPMs and Systems Biology: two complementary theoretical approaches to developmental molecular regulatory networks

### 2.1 DPM theory links genetic and physical morphogenetic processes

Stuart Newman and co-workers postulate the concept of “Dynamical Patterning Modules” (DPMs). In line with early theories of development (Thompson, 1942; Turing, 1952), DMPs constitute a theoretical framework that couples the evolutionary, developmental and morphogenetic role of specific molecules with the generic physical and chemical processes acting on living matter (Newman & Bhat, 2008, 2009; Hernández-Hernández *et al.*, 2012) (Figure 1, right). DPMs are defined as sets of gene products and other molecules in conjunction with the physical morphogenetic and patterning processes they mobilize, in the context of multicellularity. The DPM framework provides explanations to how cell clusters develop into the characteristic morphologies of chemically and mechanically excitable mesoscopic materials (e.g., hollow, multilayered, elongated, segmented forms; Newman & Bhat, 2009). This framework has been fruitfully used to study morphogenetic principles in animal and plant development (Newman & Bhat, 2008, 2009; Zhu *et al.*, 2010; Hernández-Hernández *et al.*, 2012; Benítez *et al.*, 2018). Up until now, the DPM framework has not been used to study the development of an aggregative multicellular organism.

Newman and co-workers employed the term “module” to emphasize the semi-autonomous behavior of a DPM, but DPMs can still interact with each other, forming a large network and generating a complex “patterning code”. However, the DPM framework overlooks the regulatory networks underlying each module and how multiple modules can dynamically interact to achieve a given temporal coordination of DPM activation. We argue that the concept of “Network Motifs” from systems biology can shed further light on the dynamic properties of DPMs in terms of their intrinsic nonlinear molecular dynamics.

### 2.2. Network Motifs links Molecular Regulatory Networks and generic dynamical processes

Systems biology arose during the last century as an interdisciplinary field aiming to understand, from a mathematical and theoretical perspective, the complexity of living systems and, in particular, the molecular regulatory networks (MRNs). Systems biology perceives complex cell functions as regulated nonlinear dynamical processes, not as the product of a static mapping from genotypes to phenotypes (Monod & Jacob, 1961). One of the key concepts of system biology is that of Network Motifs (Alon, 2007), which helps to understand complex cell functions, such as the robust maintenance of periodic cellular cycles (Hardin *et al.*, 1990; Skotheim *et al.*, 2008) and cellular decision making (Xiong & Ferrell, 2003; MacArthur *et al.*, 2009), in terms of simple and generic feedforward and feedback motifs (Figure 1, left). These motifs provide a dictionary of prototypical nonlinear dynamical behaviors (Milo *et al.*, 2002) that are shared across lineages. While the same motif can result in different biological functions in different systems, its potential nonlinear dynamical behaviors are fully determined by its topology. The concept of Network motifs has been successfully used to dissect and understand developmental MRNs (see Davidson & Levin, 2005; Prill *et al.*, 2005, for representative examples). However, the morphogenetic aspect of development is often ignored in those works and a “network motif theory of morphogenesis” remains to be consolidated.

Synthetic biology has been fundamental in testing and developing systems biology principles, showing that generic molecular behaviors can be implemented by designing the associated nonlinear dynamical behavior into network motifs (Elowitz & Leibler, 2000; Pomerening *et al.*, 2003; Prochazka *et al.*, 2017). Albeit the main focus of classical systems and synthetic biology has been on a single-cell level (*e.g.* Del Vecchio & Murray, 2015), recent studies have designed synthetic MRNs to temporally and spatially regulate the expression of genes with a morphogenetic function and generate desired multicellular structures (Toda *et al.*, 2018; Glass & Riedel-Kruse, 2018). Interestingly, the resulting “morphogenetic toolboxes” resemble synthetic DPMs. The success of these recent approaches further suggests a general synergy between systems biology and DPM theory in understanding and designing developmental and evolutionary processes.

## 3. Dissection of *M. xanthus* developmental MRN uncovers the key role of network motifs for DPM coordination

Analogous to what has been done for plant and animal systems, we postulate a set of DPMs involved in *M. xanthus* development (Table 1). Taking the MRN associated with the multicellular development of this bacterium (Arias del Angel *et al.*, 2018), we carry out a functional classification of the network nodes based solely on their reported biological role and assign them to the corresponding DPM (Table1, Figure 2 and methods). Interestingly, when incorporating the functional classification to the regulatory network we notice that the topology of each DPM is that of well known network motifs. Thus, we propose a new DPM structure, classifying its components into three categories: (1) *Patterning Nodes* are those that have an experimentally annotated developmental and/or morphogenetic function and thus determine the identity of the DPM they belong to; (2) *Motif Nodes* are the ones that create dynamical motifs (in the sense of Systems Biology) through their interactions with Patterning Nodes; and (3) *Coupling Nodes* are those that connect two or more DPMs and coordinate their dynamics.

**Table 1.**
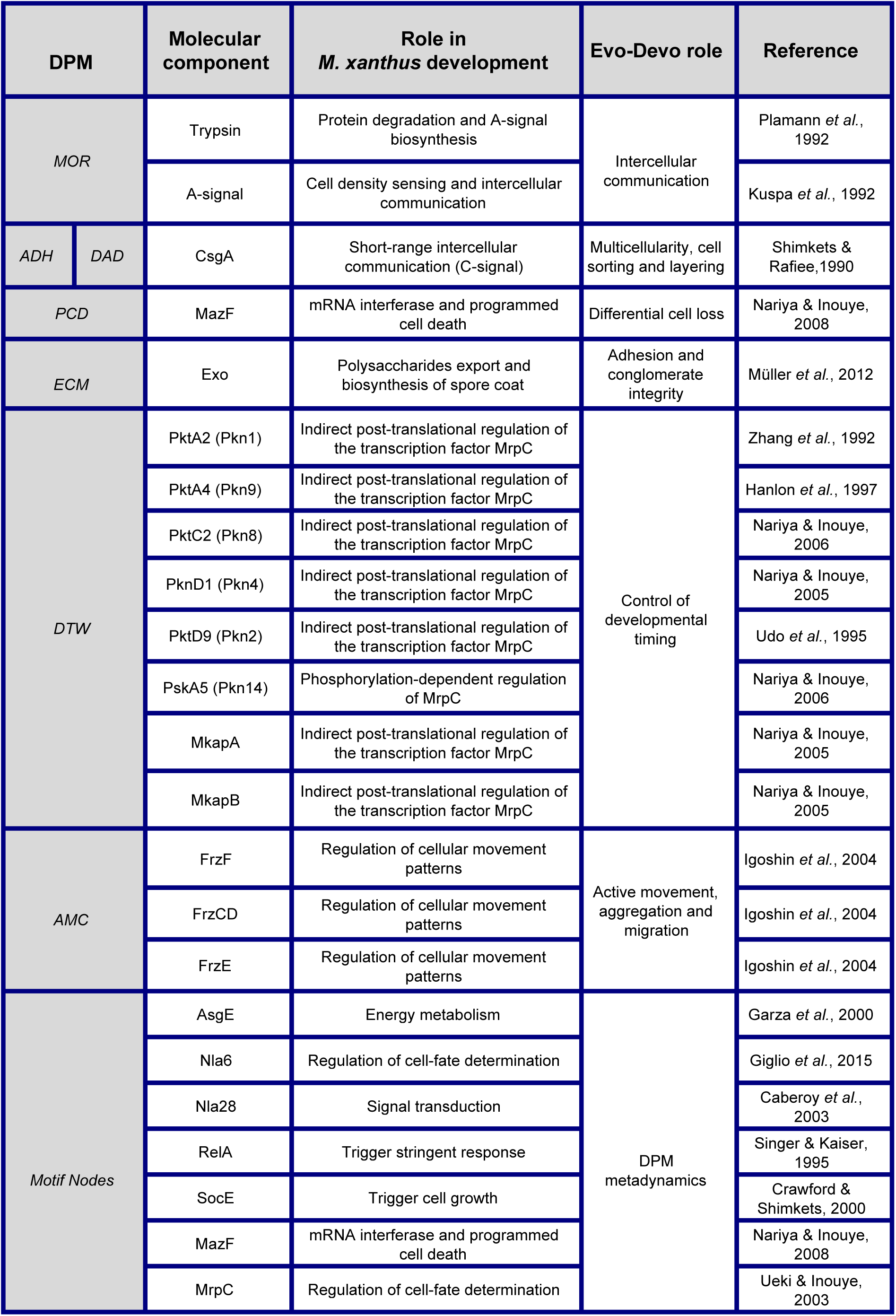
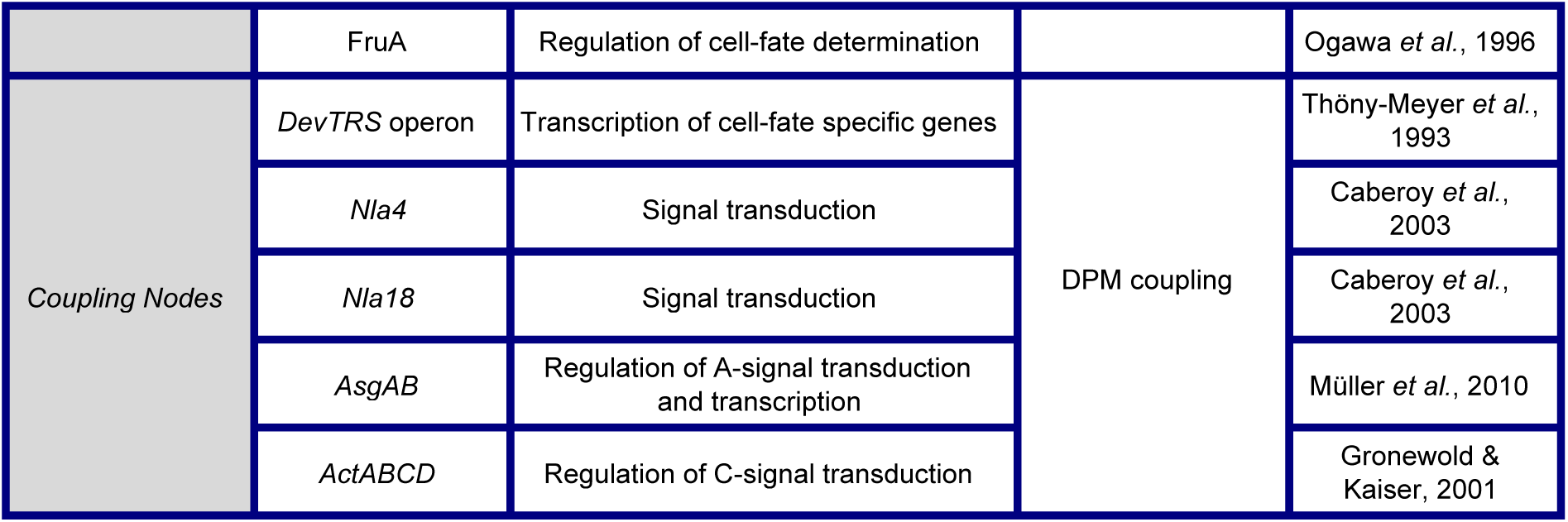
DPM assignment to molecular components involved in *M. xanthus* multicellular development. DPMs: morphogen (*MOR*), adhesion (*ADH*), differential adhesion (*DAD*), programmed cell death (*PCD*), extracellular matrix (*ECM*), developmental time window (D*TW*), active movement coordination (*AMC*), Motif Nodes and Coupling Nodes.

**Figure 2.**
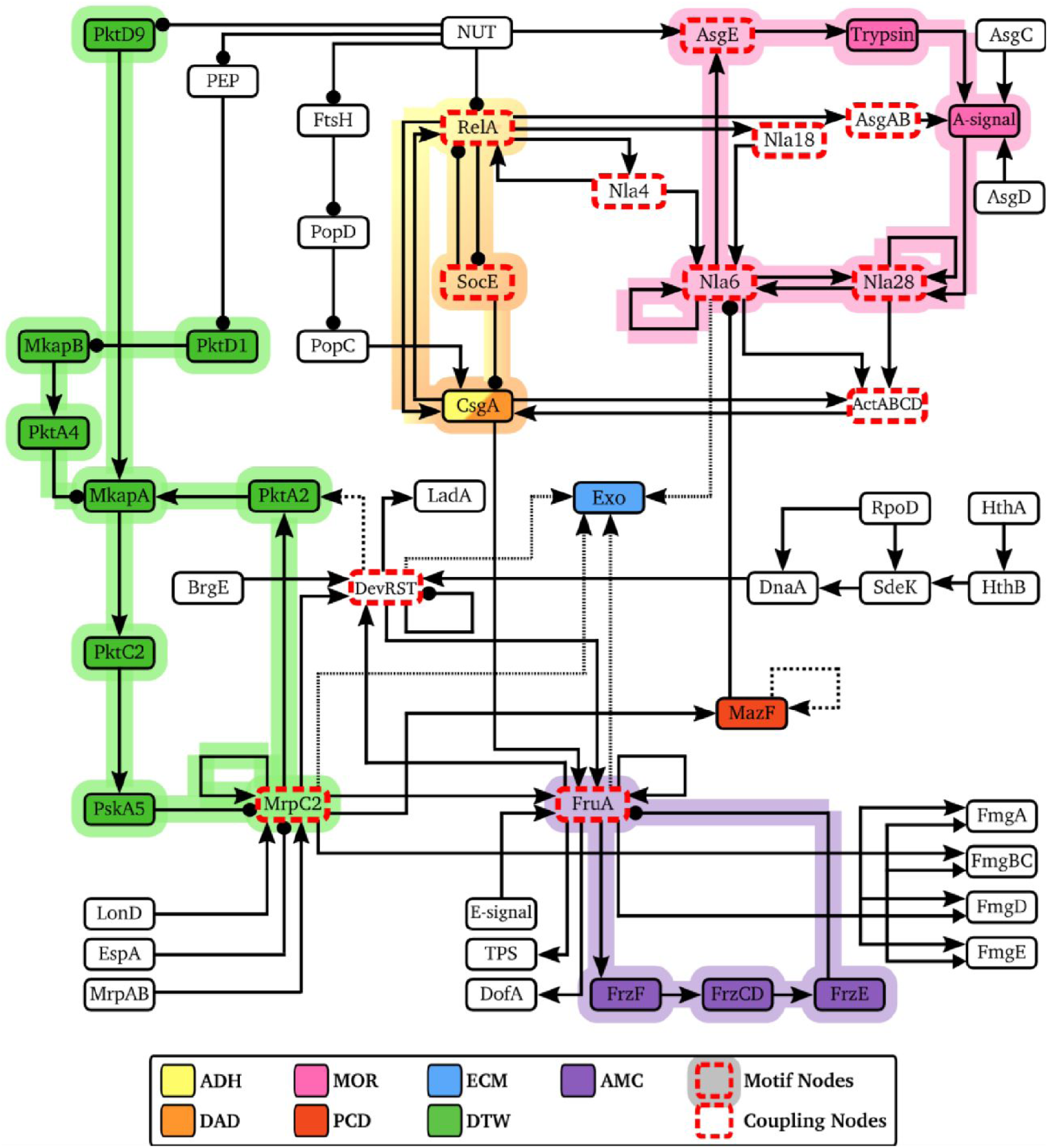
*M. xanthus* developmental regulatory network and identified DPMs. The DPMs involved in *M. xanthus* multicellular development are highlighted in different colors. Each DPM is composed by Patterning Nodes (marked with the corresponding solid color) and Motif Nodes (marked with a red dashed outline and highlighted in the corresponding color). Finally, Coupling Nodes (marked with a red dashed outline) connect two or more DPMs and coordinate their dynamics. Note that Motif Nodes are also Coupling Nodes in some instances. ADH: Adhesion DPM; DAD: Differential Adhesion DPM; MOR: Morphogen DPM; ECM: Extracellular Matrix DPM; PCD: Programmed Cell Death DPM; DTW: Developmental Time Window DPM; AMC: Active Movement Coordination DPM. Arrow heads mean positive interactions (activation). Circles mean negative interactions (inhibition). Solid lines represent experimentally annotated interactions. Dashed lines represent computationally predicted interactions. (Developmental MRN taken from Arias del Angel *et al.*, 2018)

We define one DPM as the subnetwork of associated Patterning and Motif Nodes. We hereby list the identified nodes that compose each DPM in *M. xanthus* (section 3.1) and analyze their temporal behaviors based on the network motifs that comprise each DPM (section 3.2 and 3.3).

### 3.1 Identification of DPMs in *M. xanthus*: patterning, motif and coupling nodes

#### Adhesion (ADH) and Differential Adhesion (DAD)

Adhesion (ADH) and differential adhesion (DAD) are the first and best described DPMs, both in animal and plant development (Newman & Bhat, 2009; Hernández-Hernández *et al.*, 2012), given that adhesion between neighboring cells is an essential property of multicellularity (Abedin & King, 2008). We identified a single Patterning Node belonging to ADH and DAD, CsgA, a polarized membrane protein that is involved in adhesion and intercellular communication (Lobedanz & Søgaard-Andersen, 2003). Two Motif Nodes were identified forming a direct (via RelA) and an indirect (via SocE) positive feedback loop with CsgA (Arias Del Angel *et al.*, 2018). This motif is also known as bistable or toggle switch as it is the basic motif for bistability, the coexistence of two possible stable equilibrium states corresponding to robust cell decisions (Ferrell & Machleder, 1998; Gardner *et al.*, 2000).

#### Morphogen (MOR)

Diffusive molecules allow intercellular communication and affect cell identity and/or behavior in a concentration dependent manner (Turing, 1952; Green & Sharpe, 2015). Concentration gradients that emerge from passive or active mechanisms determine the way in which diffusive molecules create ordered spatial structures in multicellular organisms (“morphogenesis”, Turing, 1952). Two Patterning Nodes from *M. xanthus* network were classified into this DPM: A-signal and Trypsin. A-signal is a cell density signal, consisting of small diffusive peptides cleaved by the protease Trypsin (Kuspa *et al.*, 1992), that indicates when to start forming fruiting bodies. The MOR Motif Nodes, AsgE, Nla6 and Nla28, (Arias Del Angel *et al.*, 2018) form an indirect positive feedback loop with MOR Patterning Nodes, creating a toggle switch and thus bistability and robust cell decision-making.

#### Extracellular Matrix (ECM)

The extracellular matrix is a complex macromolecular network that surrounds cell aggregates and forms part of their microenvironment. ECM composition can drastically vary through time and between different aggregates, determining their viscoelasticity and cohesivity. The ECM maintains cells together even in the absence of direct cell adhesion and it can influence the shape, pattern, polarity and behavior of multicellular structures (Daley *et al.*, 2008; Theocharis *et al.*, 2016; Mouw *et al.*, 2017). The Patterning Node for this DPM is the Exo locus, which contains *M. xanthus* genes that export polysaccharides to the cell surface to coat the spores within the fruiting body and allow them to stick to each other (Müller *et al.*, 2012). We did not identify any motif node for this Patterning Node, probably due to the lack of experimental characterization of its transcription factors.

#### Programmed Cell Death (PCD)

Programmed cell death is a fundamental biological strategy of multicellular structures since it can change their mass, density and shape without altering the identity of the remaining cells (Jacobson *et al.*, 1997), as observed in the formation of fingers and other extremities during morphogenesis (Jacobsen *et al.*, 1996; Milligan *et al.*, 1995). It also allows the elimination of dysfunctional individuals (*e.g.* cancerous cells) and ensures survival of the population (Jacobson *et al.*, 1997). This DPM was previously named APO, for apoptosis (Newman & Bhat, 2008, 2009), but we rename it here to account for different cell death mechanisms. MazF is an endonuclease that is implicated in programmed cell death during fruiting body development of some *M. xanthus* strains (Nariya & Inouye, 2008). Thus, it is included as a PCD Patterning Node. As in the case of ECM, we were not able to identify any Motif Nodes, nonetheless MazF itself is a Coupling Node between the MOR and DTW DPMs (see below).

All the DPM types mentioned so far were previously identified as necessary for both animal and plant development (Newman & Bhat 2008, 2009; Hernández-Hernández *et al.*, 2012). However, some fundamental genes of the *M. xanthus* developmental MRN could not be framed in any existing DPM. Therefore, we define two new DPMs based on their role during *M. xanthus* development: *Developmental Time Window (*DTW) and *Active Movement Coordination (AMC)*. Similar to the Oscillation (OSC) DPM (described in Newman & Bhat, 2008), it is the temporal, more than the stationary, nature of these DPMs that largely determines their functions.

#### Developmental Time Window (DTW)

The function of this DPM is to temporally regulate the onset of developmental processes. It is responsible for responding to environmental cues in a timely fashion to create a specific time window during which the other DPMs can be activated in a coordinated manner. This coordination initiates a developmental morphogenetic process, thus affecting the final organization and patterning of the resulting multicellular structure. In *M. xanthus*, this DPM responds to the lack of nutrients in the environment by creating a developmental time window that is critical for the formation of fruiting bodies with the proper morphology and spatial organization. It has indeed been shown that knocking out various DTW Patterning Nodes alters the onset of fruiting body formation after sensing the starvation signal, affecting fruiting body size, spore number and viability, and the spatial pattern exhibited by the fruiting body population (Escalante *et al.*, 2012; Rivera-Yoshida *et al.*, 2019). All the Patterning Nodes in this DPM are post-translational regulators of the transcription factor MrpC2, which is involved in the cell fate decision-making during development (Nariya & Inouye, 2006). MrpC2 is classified as a Motif Node since it forms two well-known motifs that reflect the timing role of DTW: an incoherent feedforward loop (IFF; Milo *et al.*, 2002) and an excitable fast-positive/slow-negative feedback loop (Izhikevich & FitzHugh, 2006; Tsai *et al.*, 2008, Franci *et al.*, 2018). The IFF motif involves the input cascades from NUT (*i.e.* nutrient levels in the environment sensed by the cells) to MkapA passing through PktD1 and PktD9, two parallel paths with an overall opposite sign. The excitable fast-positive/slow-negative motif involves the nodes in the feedback loop closed around Mrpc2 (the Motif Node). In the next section, we describe in detail the coupled dynamics of these two motifs.

#### Active Movement Coordination (AMC)

This DPM regulates and coordinates the spatial direction of active motion of single cells. It can induce a periodic reversal in the direction of motion by switching the polarity of the molecular motor in *M. xanthus*. Instead of a morphogenetic gradient, it is the synchronization of this oscillatory movement that determines the final spatial distribution of cells and, in turn, fruiting bodies (Sager & Kaiser, 1994; Igoshin *et al.*, 2004). When knocking out any of the Patterning Nodes of this DPM (FrzCD, FrzE and FrzF), cells fail to synchronize their periodic movement and to correctly form fruiting bodies (Igoshin *et al.*, 2004). The three Patterning Nodes in AMC, together with their Motif Node (FruA), form the same fast-positive/slow-negative feedback loop observed in DTW. This motif can produce sustained oscillations, which are responsible for alternation of movement behavior as suggested by Igoshin *et al.*, 2004 and as detailed in the next section.

### 3.2 The rich dynamics of DPMs

Beside their semi-autonomous morphogenetic role, DPMs are embedded in complex MRNs and participate in the rich dynamics of these networks. A key observation of our dissection of the *M. xanthus* developmental MRN is that most DPMs possess a clear network motif structure. The network motif structure of each DPM determines its endogenous dynamical behavior, its dynamical response to exogenous inputs, and its dynamical interaction with other DPMs. The function of a DPM is thus mainly determined by its morphogenetic role, whereas its temporal behavior is mainly determined by its motif structure. The three motifs we identified are: a bistable and multistable switch (in ADH/DAD and MOR); a fast-positive/slow-negative feedback loop (in DTW and AMC); and an incoherent feedforward loop (IFF) (in DTW). Except for the latter, these motifs involve at least one Motif Node, i.e., a node without an annotated morphogenetic function in the classical sense of the DPM theory. Motif Nodes, and their connections to Patterning Nodes, are crucial to generate and maintain a robust nonlinear DPM dynamical behavior. Removing these nodes would affect the functioning of the DPM by disrupting its endogenous and exogenous dynamical behavior inside the network. (Figures 3 and 4). In this section we describe the dynamical behavior of each identified network motif when disconnected from the rest of the MRN and under parametric or topological perturbations.

**Figure 3.**
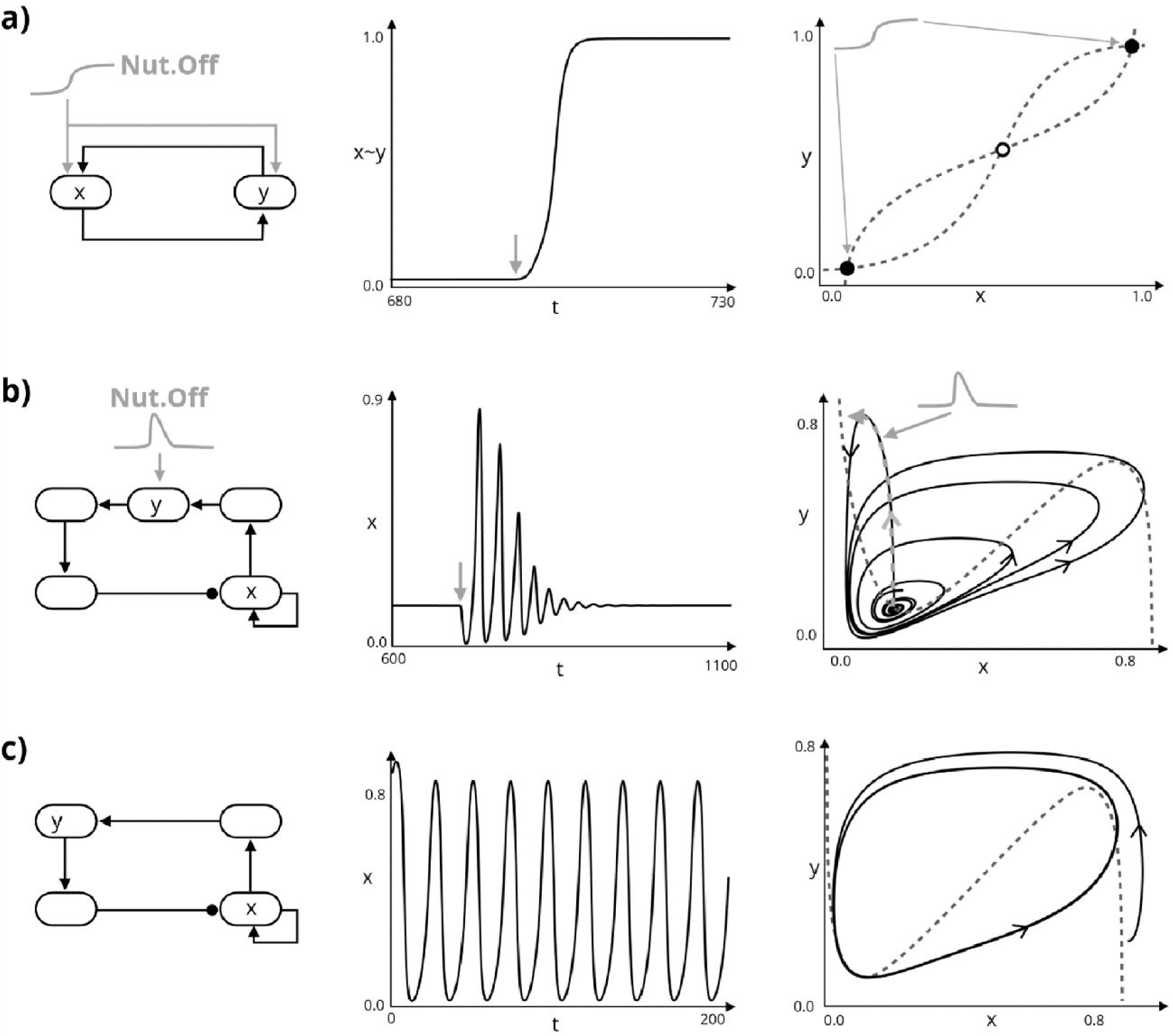
Identified network motif dynamics. a) Bistable switch motif and dynamics. The positive feedback mediated by mutual excitation underlies the binary (switch-like) response of this motif to incoming inputs (light gray). The nullcline geometry explains this behavior in the x-y phase portrait (right), where the unstable equilibrium is drawn as a white circle and the two stable equilibria as black circles. b) Fast-positive, slow-negative feedback motif and its dynamics in the excitable regime. The fast positive feedback underlies the switch-like response of this motif on short timescales. The long and thus slow negative feedback transforms the fast switch-like behavior into a transient excitation that can be understood as a slow adaptation around the fast bistable nullcline (right). c) Fast-positive, slow-negative feedback motif and its dynamics in the oscillatory regime. In the presence of a constant excitation, or in the absence of a constant inhibition, the excitable behavior of this motif is turned into a sustained oscillatory behavior that can be understood as a periodic slow adaptation around the fast bistable nullcline (right). Inputs are indicated in light gray. Nullclines in the right plots are indicated in dashed dark gray. In b) and c) only the x-nullcline is drawn. In the motif network graphs (left) pointed arrows indicate excitation and rounded arrows inhibition. Units for x,y, and t are arbitrary. The variables x and y indicate the activity of the respective nodes in the motif graph on the left.

**Figure 4.**
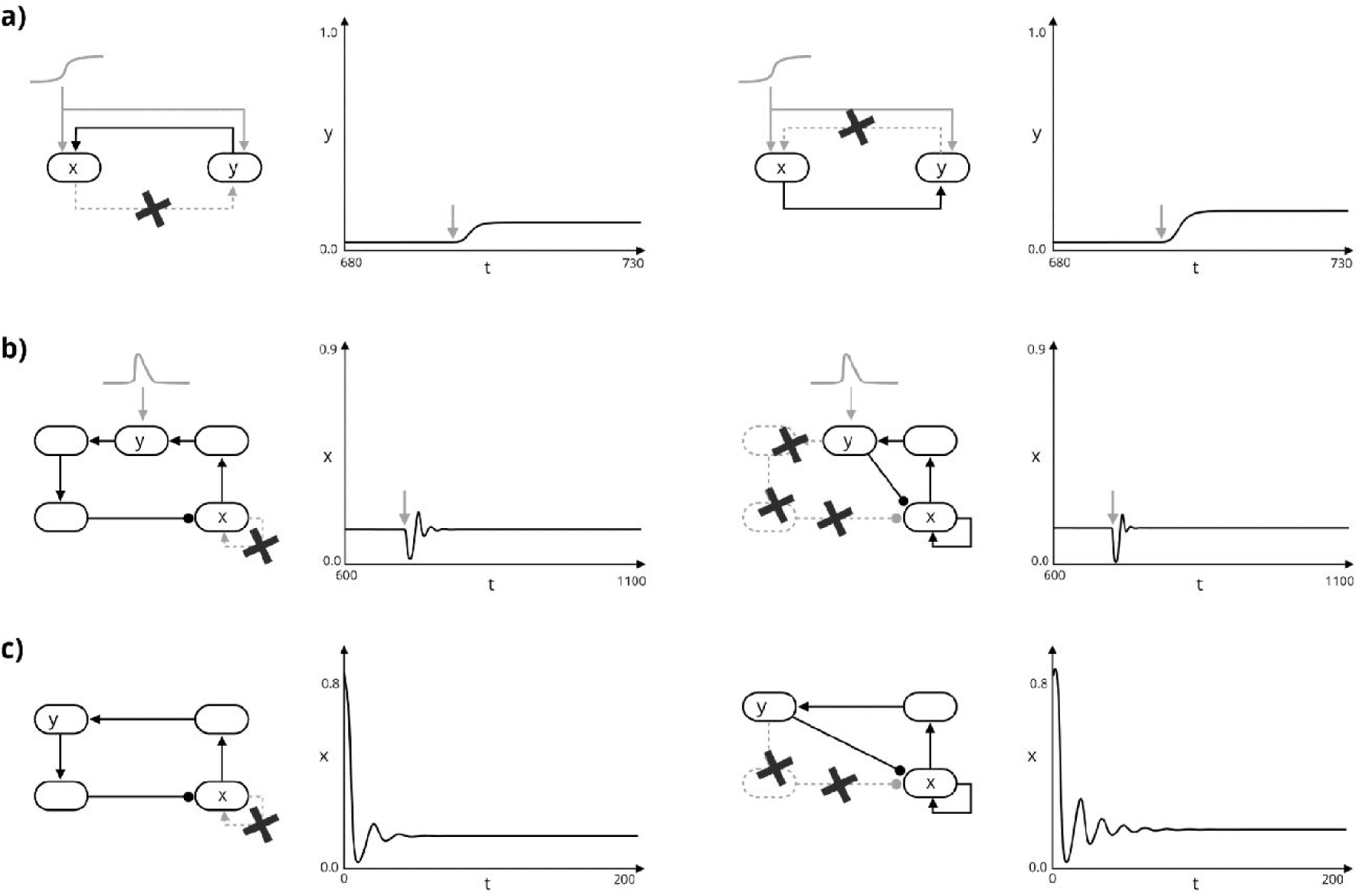
Perturbed dynamics of the identified network motifs. a) Removing mutual excitation and thus positive feedback abolishes the robust binary response shown in Figure 3a. b,c) Removing the fast positive feedback or making the negative feedback shorter (and thus faster) abolishes the excitable (b) and the sustained oscillatory (c) behaviors, shown in Figure 3b and 3c, respectively. Inputs are indicated in light gray. Units for x,y, and t are arbitrary. The variables x and y indicate the activity of the respective nodes in the motif graph on the left.

In the ADH/DAD and MOR DPMs we identify multiple variations of a bistable or multistable switch, a motif that is characterized by positive feedback (Figure 3a) arising from either mutual excitation or mutual inhibition (known as “toggle switch”; see section 5.3 of Del Vecchio & Murray, 2015). Bistability engenders discrete-like transitions in response to continuous inputs. In bistable systems, small inputs push the system towards a low state, regardless of their exact magnitude, whereas large inputs push the system towards a high state, regardless of their exact magnitude (Figure 3a, central plot). Intermediate-sized inputs can either push the system toward a high or low state, depending on the system’s state when the input is received, meaning that a bistable switch has memory. Removing the positive feedback from a bistable switch, for instance by eliminating or reducing mutual excitation, results in the system responding linearly to incoming signals (Figure 4a). This linear response is insufficient to maintain memory of the received inputs and to robustly broadcast them to the rest of the network. Hence, it is not surprising that bistability is crucial for robust cell regulation and decision making (Xiong & Ferrell, 2003; MacArthur *et al.*, 2009). The positive feedback loops present in the ADH/DAD DPM induce large, persistent, binary-like responses to the incoming developmental signal (*i.e.* lack of nutrients in the medium), thus robustly triggering *M. xanthus* developmental morphogenesis.

In the DTW DPM we find two different motifs, an IFF loop that affects a fast-positive/slow-negative feedback loop (Figure 2) as described in section 3.1. The hallmark of the first motif, IFF loop, is a linear but transient response to incoming inputs. In the DTW DPM we observe this as follows: upon nutrient depletion (NUT node off), the short path quickly turns on, leading to a transient activation of MkapA (node Y in Figure 3b) that ends as soon as the long path turns off and inhibits MkapA. The transient activation of MkapA acts as an input to the fast-positive/slow-negative feedback loop, the second motif in this DPM (Figure 3b, light gray). The fast-positive loop in this motif amplifies the transient excitation by generating a switch-like, all-or-none, response (Figure 3b, right plot) of Mrpc2 (node X in Figure 3b). The amplified nonlinear response is only transient due to the presence of the slow-negative feedback loop, which brings Mrpc2 back to rest (Figure 3b, central plot). This behavior is referred to as “excitability”, a key signaling mechanism in neural systems (Izhikevich, 2007) and possibly in MRNs (Süel *et al.*, 2006; Tomlin & Axelrod, 2007). Removing the fast-positive loop or making the slow-negative loop shorter, and therefore faster, turns the excitable behavior of Mrpc2 into a linear response that is too weak to robustly broadcast its input to the rest of the network (Figure 4b).

The AMC DPM exhibits the same qualitative topology as the fast-positive/slow-negative loop in the DTW DPM (Figure 3c). However, the two DPMs differ in their inputs. Beside the transient excitation mediated by the IFF loop, DTW is under constant inhibition (through PktD9), whereas AMC does not receive any sustained inhibitory input. In the absence of sustained inhibition, excitability turns into sustained, relaxation-like oscillations (Izhikevich & FitzHugh, 2006) (Figure 3c). As opposed to the purely inhibitory oscillator described in Igoshin *et al.* 2004, we speculate that the resulting biochemical oscillator is of the relaxation type because it arises from interlinking a fast-positive and slow-negative feedback. This type of oscillators is known for exhibiting better robustness and tunability properties than purely negative feedback oscillators (Tsai *et al.*, 2008; Franci *et al.*, 2017). Similar to the DTW DPM, removing the fast-positive loop or making the negative loop shorter (and faster) destroys the sustained oscillatory behavior (Figure 4c). In this case, the function of the AMC DPM would be completely compromised, as in previously suggested knockout conditions (Igoshin *et al.*, 2004).

### 3.3 A dynamical network of DPMs

The Coupling Nodes we postulate interconnect two or more DPMs, thus generating richer network dynamics and, eventually, richer morphogenetic outputs. Figure 5a reproduces the collective dynamics of the DPM subnetwork, which includes DTW, AMC, and ADH/DAD (from Figure 2). The environmental input of this subnetwork results from nutrient depletion (*i.e.* NUT node off) turning on the ADH/DAD bistable switch and activates DTW’s excitable dynamics, as detailed in section 3.2. These initial responses are transmitted to the DPM subnetwork, which results in Mrpc2 exciting FruA, DevRST, and MazF; DevRST exciting PktA2 and FruA; and CsgA exciting FruA (see methods for the equations and the Julia Notebook in the code section for details). We analyze and simulate this subnetwork as it contains most of the nodes for which dynamical developmental roles have been experimentally annotated and for which genetic knockout experiments have been previously performed.

**Figure 5.**
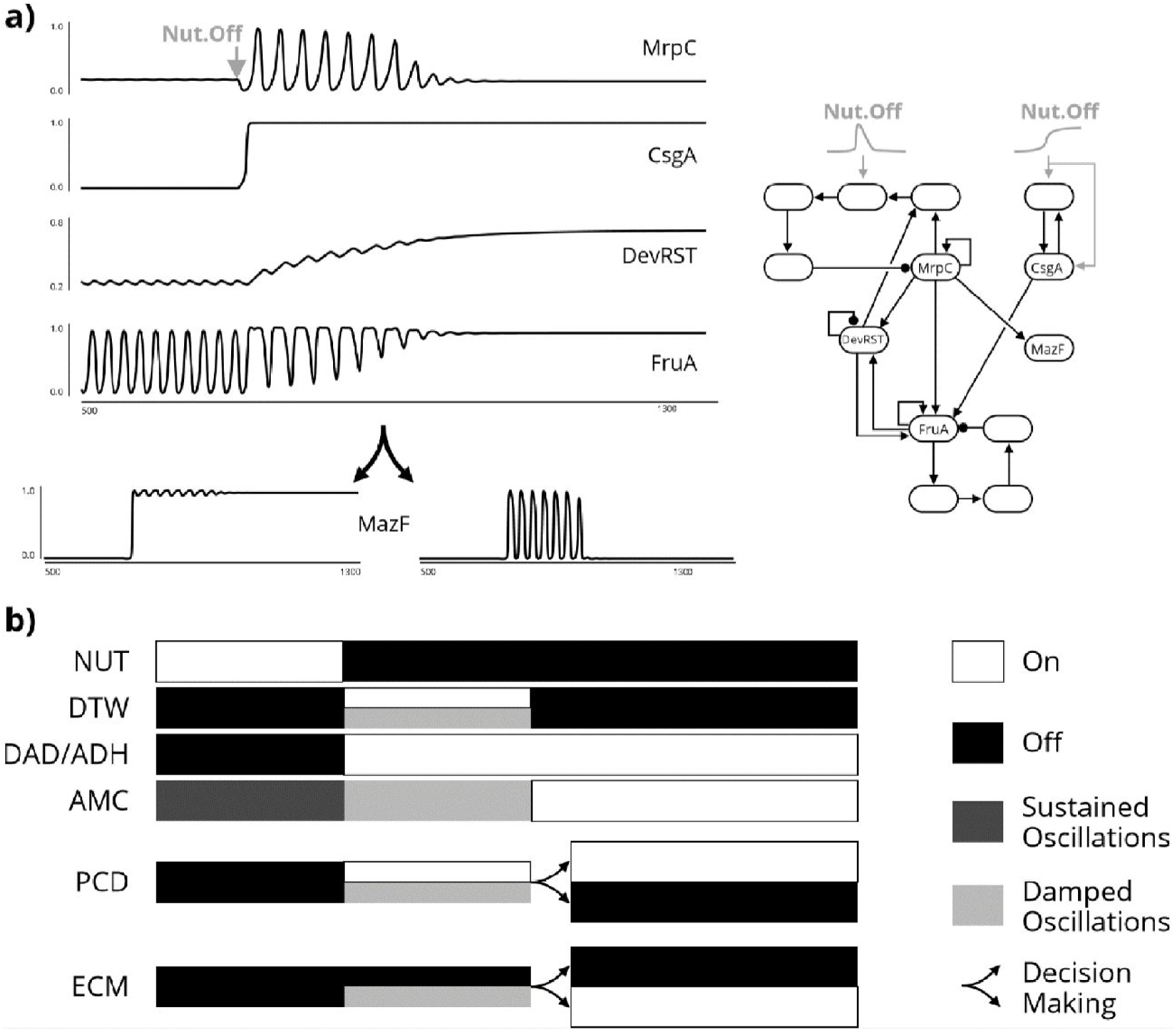
Coupled DPM dynamics predict biologically relevant developmental transition from vegetative state to either spore formation (ECM) or programmed cell death (PCD). a) Simulated network structure (right) and temporal behavior of relevant nodes (left). In the simulated network structure, the light gray “Nut Off” inputs are sketches of the time-varying inputs used to simulate the effect of nutrient depletion on the stimulated nodes. The pulse-like left trace arises from the IFF path of the DTW DPM. Units for time and simulated variables are arbitrary. b) Schematic of the simulated dynamical transitions corresponding to the associated DPM behaviors and biological interpretation (time axis and intervals are arbitrary and not on the same timescale as a)). When nutrients become scarce (the NUT node is switched from high to low), the DTW DPM bursts an excitation and the DAD/ADH DPM switches on to promote cell adhesion. This triggers a cascade of effects. The DevRST node gradually activates which induces a gradual reduction in the AMC DPM oscillation amplitude. Eventually, AMC DPM oscillations disappear, eliminating periodic cell movement reversal and favoring cell aggregation. In turn, depending on small parameter differences, the PCD DPM can either switch on permanently or only transiently. In the first case, the cell decides for its programmed death, not producing ECM proteins and becoming a spore via the ECM DPM. In the second case, the cell avoids its programmed death and activates the ECM DPM to favor spore formation.

In detail, the simulated subnetwork dynamics are the following. Nutrient depletion switches on CsgA and engages the transient activation of the DTW DPM, creating a temporal window in which rich dynamical phenomena happen: (1) the oscillation of the AMC DPM slowly turns off (thus turning off periodic cell movement reversal and favoring aggregation) under the joint effect of DevRST and CsgA; (2) DevRST slowly transitions from inactive to active, which could allow the system to transition back to the vegetative state if nutrients are restored in this timespan or if other environmental conditions change; (3) finally, the PCD DPM (node MazF) can permanently turn on or not after the transient activation of the DTW DPM, depending on slight differences in its basal expression levels (*i.e.* MazF expression levels in the absence of network interactions). In a multicellular context, these small differences could be spatially coordinated by intercellular communication (not modeled here). As opposed to other modeling frameworks, such as boolean network models (Arias Del Angel *et al.*, 2018), here it is the continuous variation of biologically controlled parameters, and not the MRN initial conditions, that determines the cell fate in a multi-attractor setting. We summarize this information in Figure 5b and push further the biological implication of our analysis by speculating the response of the ECM DPM to the PCD DPM dynamical behavior (spore formation vs. cell death). The PCD DPM indirectly inhibits the ECM DPM via Nla6. In line with the biological role of these DPMs (section 3.1), activation of the PCD DPM would disfavor activation of the ECM DPM and, in turn, inactivation of the PCD DPM would favor activation of the ECM DPM. In other words, a cell that has entered development can either commit to programmed cell death or become a spore. Finally, our simplified continuous dynamical model further reproduces the effect of PtkD9 knockout on accelerating the onset of the multicellular developmental process (Escalante *et al.*, 2012) (Supplementary Figure S1). Although explorative, the described dynamical behavior suggests a plausible connection between DPMs’ nonlinear dynamics and the temporal coordination of developmental stages.

Although simplified, our model for the DPM subnetwork reproduces the following experimental evidence. First, the transient activation of MrpC2, the transcription factor and Motif Node of the DTW DPM (Lee *et al.*, 2012). Second, the developmental decision-making role of MazF, a key mediator of programmed cell death (Nariya & Inouye, 2008) and the only node of the PCD DPM. Third, our model agrees with previous modelling efforts that suggest a transition of FruA (belonging to the AMC DPM) post-translational dynamical behavior from oscillatory to equilibrium (Igoshin *et al.*, 2004). The proposed numerical simulations are exploratory. Nonetheless, the simulated dynamics of the three nodes MrpC2, MazF, FruA (for which experimental data exist, see above) qualitatively fit the experimentally measured temporal variations during development. The dynamics of the remaining nodes is speculative but suggestive of the following connection between DPM activation and developmental stages (Figure 5 and legend for details).

## 4. Discussion and outlook

In line with what has been previously proposed for plant and animal systems, we have postulated a set of DPMs associated to the developmental behavior of *M. xanthus*, which, as an aggregative multicellular microorganism, opens new avenues for wide comparative analyses among multicellular lineages. In this study we identified and analyzed a subset of DPMs that are common to most plants, animals and this multicellular bacterium: Adhesion (ADH), Differential Adhesion (DAD), Morphogen (MOR), Extracellular Matrix (ECM) and Programmed Cell Death (PCD, or APO in previous work) (Benítez *et al.*, 2018; Newman & Bhat, 2008, 2009). We also identified two new DPMs that are important for multicellular development: Developmental Time Window (DTW) and Active Movement Coordination (AMC). It is worth noting that in this case, and in contrast to what has been previously done for plants and animals, the DPMs have been inferred from a single model organism. Therefore, it would be important to confirm the occurrence of these DPMs in other multicellular systems, such as other eukaryotic aggregative organisms (*e.g.* dictyostelids), synthetic multicellular aggregates, as well as animals and plants.

While the DPM framework is now well-established and has led to the detailed characterization of specific and semi-autonomous DPMs (e.g. Zhu *et al.*, 2010; Hernández-Hernández *et al.*, 2018), little is known about the dynamics of these modules, how they propagate environmental signals, and how they connect to each other in developmental processes. Besides characterizing the DPMs for aggregative bacteria, one of the objectives of this work was to take Network Motifs tools and concepts to explore the dynamics of the molecular elements associated with each DPM and their connections. In turn, this would enable a much-needed connection between the molecular scale of systems biology and the organismic scale of morphogenetic processes. This approach revealed that some of the dynamic motifs and mechanisms that might, at least partially, underlie the robust behavior, dynamic abundance and semi-autonomy of DPMs. For instance, the AMC DPM involving a relaxation-like oscillator, interlinking fast-positive and slow-negative feedbacks, exhibits better robustness and tunability properties than a purely negative feedback oscillator.

Our analysis leads to the classification of what we call Motif Nodes, underlying DPM nonlinear dynamics, and Coupling Nodes, underlying DPM temporal coordination. Motif Nodes allowed us to identify molecules that enable specific dynamical behaviors. Variation in these nodes may underlie developmental qualitative variation of potential evolutionary relevance, not because of an intrinsic property but because of their role as part of complex motifs and DPMs. Similarly, variations in Coupling Nodes may underlie significant developmental, and thus morphological, changes. Recent studies have demonstrated that the topology of a network can result in different dynamical behaviors depending on environmental conditions and network interaction strengths (Jimenez *et al.*, 2017; Perez-Carrasco *et al.*, 2018; Verd *et al.*, 2019). In *M. xanthus* developmental MRN, Motif and Coupling Nodes have an important role in generating various DPM dynamics that trigger and drive multicellular development after nutrient depletion. The resulting dynamical behaviors are determined by the specific motif topology and can vary depending on control inputs, parameters, and initial conditions. This controlled variability in DPM dynamics supports the idea of diverse multicellular morphologies developing and evolving through the dynamic interaction between multiple DPMs and the environment.

The functional classification of network nodes into Patterning, Motif and Coupling Nodes may be particularly suitable to pursue broad comparative studies, in which analyses that are solely based on the molecular or phylogenetic nature of the nodes can be obscured by the evolutionary molecular drift (*e.g.* Arias Del Angel *et al.*, 2017). It is indeed the generic (physical and dynamical) properties of developmental MRNs’ constituents, rather than their genetic or molecular characterization, that might enable novel and more robust Evo-Devo comparative studies. This idea is already present in DPM theory but only regarding physical aspects of development and evolution, while largely neglecting dynamical and purely temporal ones. Considering the two together, as suggested by our work, reveals that physical and temporal aspects of development are tightly intertwined at the transition to multicellularity. The fact that many DPMs, with generic morphogenetic functions, are comprised by highly recurrent network motifs, with generic dynamical behaviors, suggests that evolution is not only constrained by the physical properties of the materials on which it acts upon, but also (and equally important) by the possible dynamical behaviors that the emerging molecular networks are likely to exhibit, as dictated by the topology of simple (and thus highly recurrent) network motifs.

We conclude that the integration of the DPM framework and Network Motif theory can lead to a more comprehensive understanding of the processes underlying multicellular development and morphogenesis, gathering knowledge from molecular networks at different scales (from single DPMs to the organismic level). In this case, our approach allowed us to uncover some of the dynamic features behind the aggregative development of myxobacterial fruiting bodies, thus contributing to a better understanding of both the specific and common aspects of the different paths to multicellularity.

## Methods

### DPM identification

Analysis of *M. xanthus* developmental MRN (taken from Arias del Angel *et al.*, 2018) started with a more detailed functional classification of each component. The classification was based only on their biological functions and the process they are involved in. This information was extracted from the literature found for each node (Table 1) and from the Gene Ontology database (Ashburner *et al*., 2000, The Gene Ontology Consortium, 2019). Based solely on the initial functional classification, we assigned the corresponding DPM to each node (Table 1). After this second classification we incorporated the regulatory interactions and found that the nodes belonging to the same DPM form a dynamic structure. Patterning Nodes group together and form well-known network motifs with at least one Motif Node, these “groups” are connected via Coupling Nodes (N.B. some nodes can be both Motif and Coupling Nodes). Once we identified these DPM structures, we analyzed their temporal behaviors through computational simulations.

### Model equations and numerical simulations

The model equations are generic in the sense that they capture the generic properties of MRN (sigmoidal activations, linear degradations, feedback loops) in simplified equations that do not necessarily describe the biomolecular details. As such, they are amenable to a mechanistic mathematical analysis, as provided in the text, without compromising the qualitative value of the predictions. These equations can be found in supplementary information.

The parameters used for our simulations are based on experimental evidence reported in the literature for the transient activation of MrpC2 (Lee *et al.*, 2012) (DTW), the decision-making role of MazF (Nariya & Inouye, 2009) (PCD), the knockout effect of PtkD9 knockout (Escalante *et al.*, 2012), and the oscillatory behavior of FruA and the Frz genes (AMC) previously proposed (Igoshin *et al.*, 2004).

We use our model to explore perturbations to the network motifs and to explore the simplified DPM subnetwork, predicting dynamic coupled behaviors (on, off and sustained or damped oscillations) for each DPM that replicate the relevant transitions during *M. xanthus* multicellular development.

### Code availability

The code and equations used to generate Figure 3 and 4 are available at https://github.com/NeurAlessio/Myxo-DPM-SB/blob/master/TIM-MOV-SWI-lmcbIsolated.ipynb

The code used to generate Figure 5 is available at https://github.com/NeurAlessio/Myxo-DPM-SB/blob/master/TIM-MOV-SWI.ipynb

The code used to generate Figure S1 is available at https://github.com/NeurAlessio/Myxo-DPM-SB/blob/master/TIM-MOV-SWI-DeltaPktD9.ipynb

## Acknowledgements

During the peer review process, Juan A. Arias Del Angel, our dear friend and coauthor, passed away. He was intimately involved in the analyses, discussions and writing the manuscript. We dedicate this paper to his memory.

The authors would like to thank Stuart Newman for valuable feedback on the manuscript. A.G.H. is currently a doctoral student at University College London (UCL) and is funded by University College London (UCL) through an Overseas Research Scholarship and a Graduate Research Scholarship; and by Consejo Nacional de Ciencia y Tecnología (CONACyT) [scholarship number 471963]. J.A.D.A. was a doctoral student from Programa de Doctorado en Ciencias Biomédicas, Universidad Nacional Autónoma de México (UNAM) funded by CONACyT [scholarship number 580236]. N.R.Y. is funded by Programa de Becas Posdoctorales en la UNAM, A.F. and M.B. are funded by UNAM-DGAPA-PAPIIT, grant IA105518. AF is also funded by CONACyT, grant A1-S-10610.

## Supplementary material

**Figure S1.**
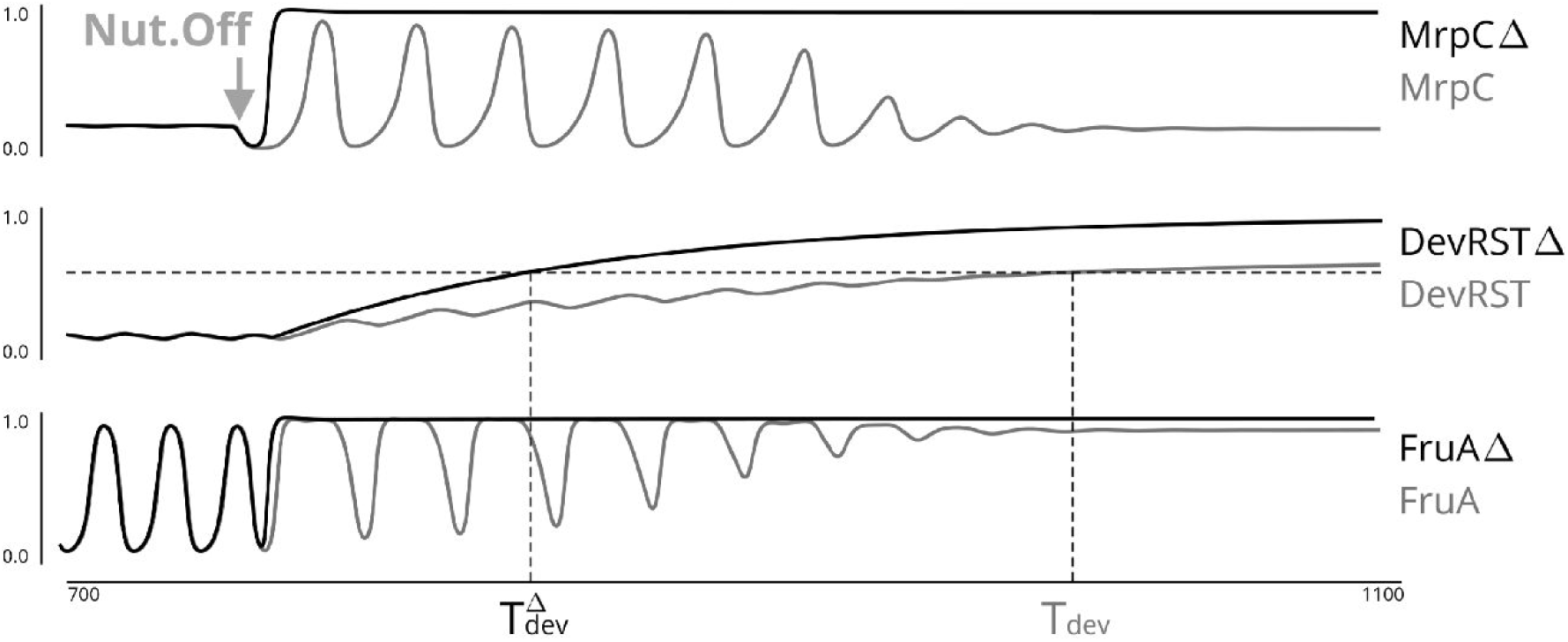
Effect of PktD9 knockout in the coupled DPM network. In light grey is the temporal behavior of relevant nodes (from Figure 5a). In dark grey is the predicted effect of knocking out each node, which replicates the experimentally effect on the accelerated onset of *M. xanthus* development (Escalante *et al.*, 2012; Arias Del Angel *et al.*, 2019). T_dev_ indicates development time in control conditions and T_dev_^Δ^ indicates development time in knock-out conditions.

### Methods and model

In the equations, 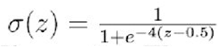.

**Equations for Figure 3a.** The used equations describe a generic two-dimensional bistable switch with an exogenous input modeling the effect of the NUT node turning off:

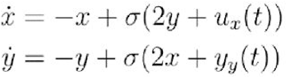

where *u*_*x*_*(t)* = 0.2(tanh(*t* - 700) + 1) - 0.6, *u*_*y*_*(t)* = 0.2(tanh(*t* - 700) + 1) - 0.4.

**Equations for Figure 3b.** The used equation s describe a generic fast-positive/slow-negative feedback motif with an instantaneous autocatalysis (positive feedback) loop and a slow, four-node negative feed back loop, under the effect of an exogenous input modeling the effect of the NUT node turning off:

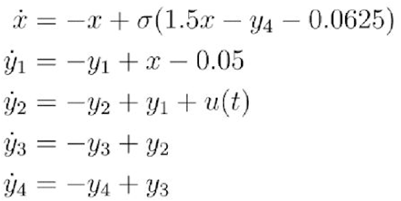

wher *u(t)* = 0.5(tanh(*t* - 700) + 1) - 0.5(tanh (0.5(*t* - 705)) + 1). In the figure, *y* = *y*_2_.

**Equations for Figure 3c.** The used equations describe a generic fast-positivc/slow-negative feedback motif with an instantaneous autocatalysis (positive feedback) loop and a slow, three-node negative feedback loop

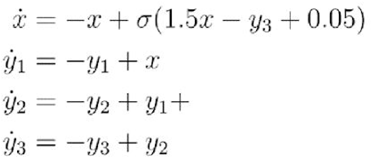

In th figure, *y* = *y*_2_.

**Equations for Figure 4a.** Same equations as Figure 3a, but with either of the mu tual excitation branches removed. Figure 4a, left:

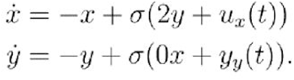

Figure 4a, right:

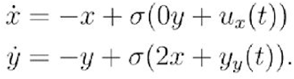

**Equations for Figure 4b left.** Same equations as Figure 3b, but with the auto-catalysis branch removed:

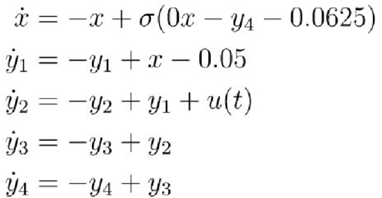

**Equations for Figure 4b right.** Same equations as Figure 3b, but with two instead than four nodes in the slow negative feedback loop:

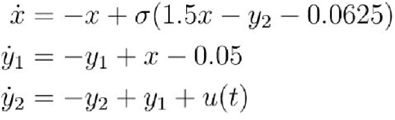

**Equations for Figure 4c left.** Same equations as Figure 3c, but with the auto-catalysis branch removed:

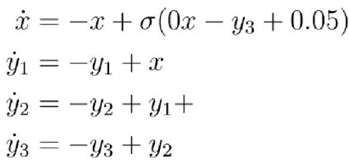

**Equations for Figure 4c right.** Same equations as Figure 3c, but with two instead than three nodes in the slow negative feedback loop:

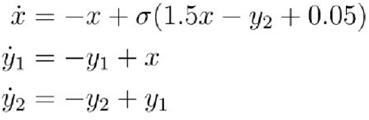

**Equations for Figure 5.**

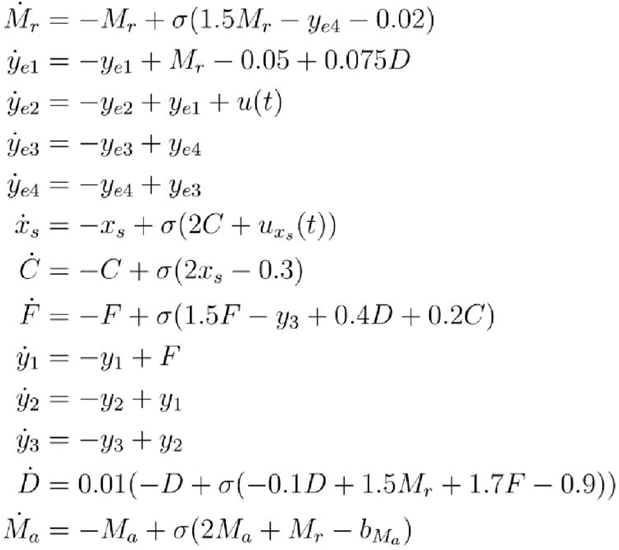

where

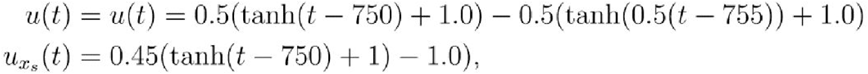

and 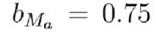 for switch-on case and 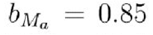 for transient case. In the figure, MrpC = *M*_*r*_, CsgA = *C*, FruA = *F*, DevRST = *D*, MazF = *M*_*a*_.

## References

Abedin, M., & King, N. (2008). The premetazoan ancestry of cadherins. Science, 319(5865), 946–948. doi:10.1126/science.1151084.

Alon, U. (2007). Network motifs: theory and experimental approaches. Nature Reviews Genetics, 8(6), 450–461. doi:10.1038/nrg2102

Arias Del Angel, J. A., Escalante, A. E., Martínez-Castilla, L. P., & Benítez, M. (2017). An Evo-Devo Perspective on Multicellular Development of Myxobacteria. Journal of Experimental Zoology Part B: Molecular and Developmental Evolution, 328(1-2), 165–178. doi:10.1002/jez.b.22727

Arias Del Angel, J. A., Escalante, A. E., Martínez-Castilla, L. P., & Benítez, M. (2018). Cell-fate determination in *Myxococcus xanthus* development: Network dynamics and novel predictions. Development, Growth & Differentiation, 60(2), 121–129. doi:10.1111/dgd.12424

Arias Del Angel, J. A., Rivera-Yoshida, N., Escalante, A. E., Martínez-Castilla, L. P., & Benítez, M. (2019). Role of aggregate size, multistability and communication in determining cell fate and patterning in *M. xanthus*. bioRxiv. doi: https://doi.org/10.1101/627703

Ashburner, M., Ball, C. A., Blake, J. A., Botstein, D., Butler, H., Cherry, J. M., Davis, A. P., Dolinski, K., Dwight, S. S., Eppig, J. T., Harris, M. A., Hill, D. P., Issel-Tarver, L., Kasarskis, A., Lewis, S., Matese, J. C., Richardson, J. E., Ringwald, M., Rubin, G. M., & Sherlock, G. (2000). Gene ontology: tool for the unification of biology. The Gene Ontology Consortium. Nature genetics, 25(1), 25–29. doi:10.1038/75556

Bahar, F., Pratt-Szeliga, P. C., Angus, S., Guo, J., & Welch, R. D. (2014). Describing *Myxococcus xanthus* aggregation using Ostwald ripening equations for thin liquid films. Scientific Reports, 4, 6376. doi:10.1038/srep06376

Benítez, M., Hernández-Hernández, V., Newman, S. A., & Niklas, K. J. (2018). Dynamical Patterning Modules, Biogeneric Materials, and the Evolution of Multicellular Plants. Frontiers in Plant Science, 9, 871. doi:10.3389/fpls.2018.00871

Bonner J. (2000). First Signals: The Evolution of Multicellular Development. Princeton University Press.

Bonner J. (2016). Multicellularity: Origins and Evolution. Edited by Niklas K, Newman S. The MIT Press.

Bretl, D. J., & Kirby, J. R. (2016). Molecular mechanisms of signaling in *Myxococcus xanthus* development. Journal of Molecular Biology, 428(19), 3805–3830. doi:10.1016/j.jmb.2016.07.008

Caberoy, N. B., Welch, R. D., Jakobsen, J. S., Slater, S. C., & Garza, A. G. (2003). Global mutational analysis of NtrC-like activators in *Myxococcus xanthus*: identifying activator mutants defective for motility and fruiting body development. Journal of Bacteriology, 185(20), 6083–6094. doi:10.1128/jb.185.20.6083-6094.2003

Crawford, E. W., & Shimkets, L. J. (2000). The stringent response in *Myxococcus xanthus* is regulated by SocE and the CsgA C-signaling protein. Genes & Development, 14(4), 483–492.

Daley, W. P., Peters, S. B., & Larsen, M. (2008). Extracellular matrix dynamics in development and regenerative medicine. Journal of Cell Science, 121, 255–264. doi: 10.1242/jcs.006064

Davidson, E., & Levin, M. (2005). Gene regulatory networks. Proceedings of the National Academy of Sciences, 102(14), 4935. doi:10.1073/pnas.0502024102

del Campo, J., Sieracki, M. E., Molestina, R., Keeling, P., Massana, R., & Ruiz-Trillo, I. (2014). The others: our biased perspective of eukaryotic genomes. Trends in Ecology & Evolution, 29(5), 252–259. doi:10.1016/j.tree.2014.03.006

del Vecchio, D. & Murray, R.M. (2015). Biomolecular feedback systems. Princeton, NJ: Princeton University Press.

Elowitz, M.B. & Leibler, S. (2000). A synthetic oscillatory network of transcriptional regulators. Nature, 403(6767), 335–338.

Escalante, A. E., Inouye, S., & Travisano, M. (2012). A spectrum of pleiotropic consequences in development due to changes in a regulatory pathway. PloS One, 7(8), e43413. doi:10.1371/journal.pone.0043413

Ferrell, J.E. & Machleder, E.M. (1998). The biochemical basis of an all-or-none cell fate switch in Xenopus oocytes. Science, 280(5365), 895–898. doi:10.1126/science.280.5365.895

Franci, A., Drion, G. & Sepulchre, R. (2018). Robust and tunable bursting requires slow positive feedback. Journal of Neurophysiology, 119(3), 1222–1234. doi: 10.1152/jn.00804.2017

Gardner, T.S., Cantor, C.R. & Collins, J.J. (2000). Construction of a genetic toggle switch in *Escherichia coli*. Nature, 403(6767), 339–342.

Garza, A. G., Harris, B. Z., Greenberg, B. M., & Singer, M. (2000). Control of asgE expression during growth and development of *Myxococcus xanthus*. Journal of Bacteriology, 182(23), 6622–6629. doi:10.1128/jb.182.23.6622-6629.2000.

Giglio, K. M., Zhu, C., Klunder, C., Kummer, S., & Garza, A. G. (2015). The enhancer binding protein Nla6 regulates developmental genes that are important for *Myxococcus xanthus* sporulation. Journal of Bacteriology, 197(7), 1276–1287. doi:10.1128/JB.02408-14

Glass, D.S. & Riedel-Kruse, I.H., (2018). A synthetic bacterial cell-cell adhesion toolbox for programming multicellular morphologies and patterns. Cell, 174(3), 649–658. doi:10.1016/j.cell.2018.06.041

Green, J. B., & Sharpe, J. (2015). Positional information and reaction-diffusion: two big ideas in developmental biology combine. Development, 142(7), 1203–1211. doi: 10.1242/dev.114991

Gronewold, T. M. & Kaiser, D. (2001), The act operon controls the level and time of C-signal production for *Myxococcus xanthus* development. Molecular Microbiology, 40(3), 744–756. doi:10.1046/j.1365-2958.2001.02428.x

Grosberg R. K. & Strathmann R. R. (2007). The evolution of multicellularity: a minor major transition? Annual Review of Ecology, Evolution, and Systematics, 38, 621–654. doi:10.1146/annurev.ecolsys.36.102403.114735

Hanlon, W. A., Inouye, M., & Inouye, S. (1997). Pkn9, a Ser/Thr protein kinase involved in the development of *Myxococcus xanthus*. Molecular Microbiology, 23(3), 459–471.

Hardin, P.E., Hall, J.C. & Rosbash, M. (1990). Feedback of the Drosophila period gene product on circadian cycling of its messenger RNA levels. Nature, 343(6258), 536–540. doi:10.1038/343536a0

Hernández-Hernández, V., Niklas, K. J., Newman, S. A., & Benítez, M. (2012). Dynamical patterning modules in plant development and evolution. International Journal of Developmental Biology, 56, 661–674. doi:10.1387/ijdb.120027mb

Hernández-Hernández, V., Barrio, R. A., Benítez, M., Nakayama, N., Romero-Arias, J. R., & Villarreal, C. (2018). A physico-genetic module for the polarisation of auxin efflux carriers PIN-FORMED (PIN). Physical Biology, 15(3), 036002. doi:10.1088/1478-3975/aaac99

Higgs, P.I., Hartzell, P.L., Holkenbrink, C., & Hoiczyk, E. *Myxococcus xanthus* Vegetative and developmental cell heterogeneity. In Rajagopalan, R., Sarwar, Z., Garza, A. G., & Kroos, L. (2014). Myxobacteria: genomics, cellular and molecular biology. Caister Academic Press.

Igoshin, O. A., Goldbeter, A., Kaiser, D., & Oster, G. (2004). A biochemical oscillator explains several aspects of *Myxococcus xanthus* behavior during development. Proceedings of the National Academy of Sciences, 101(44), 15760–15765. doi:10.1073/pnas.0407111101

Igoshin, O. A., Welch, R., Kaiser, D., & Oster, G. (2004). Waves and aggregation patterns in myxobacteria. Proceedings of the National Academy of Sciences, 101(12), 4256–4261. doi:10.1073/pnas.0400704101

Izhikevich, E.M. & FitzHugh, R. (2006). FitzHugh-Nagumo model. Scholarpedia, 1(9), 1349. doi:10.4249/scholarpedia.1349

Izhikevich, E.M. (2007). Dynamical systems in neuroscience. MIT press.

Jacobsen, M. D., Weil, M., & Raff, M. C. (1996). Role of Ced-3/ICE-family proteases in staurosporine-induced programmed cell death. Journal of Cell Biology, 133(5), 1041–1051.

Jacobson, M. D., Weil, M., & Raff, M. C. (1997). Programmed cell death in animal development. Cell, 88(3), 347–354. doi:10.1016/S0092-8674(00)81873-5

Jiménez, A., Cotterell, J., Munteanu, A., & Sharpe, J. (2017). A spectrum of modularity in multi-functional gene circuits. Molecular Systems Biology, 13(4), 925. doi:10.15252/msb.20167347

Kroos, L. (2007). The Bacillus and Myxococcus developmental networks and their transcriptional regulators. Annual Review of Genetics, 41, 13–39. doi:10.1146/annurev.genet.41.110306.130400

Kurosaka, S., & Kashina, A. (2008). Cell biology of embryonic migration. Birth Defects Research Part C: Embryo Today: Reviews, 84(2), 102–122. doi:10.1002/bdrc.20125

Kuspa, A., Plamann, L., & Kaiser, D. (1992). A-signalling and the cell density requirement for *Myxococcus xanthus* development. Journal of Bacteriology, 174(22), 7360–7369. doi:10.1128/jb.174.22.7360-7369.1992

Lee, B., Holkenbrink, C., Treuner-Lange, A., & Higgs, P. I. (2012). *Myxococcus xanthus* developmental cell fate production: heterogeneous accumulation of developmental regulatory proteins and reexamination of the role of MazF in developmental lysis. Journal of Bacteriology, 194(12), 3058–3068. doi:10.1128/JB.06756-11

Lobedanz, S., & Søgaard-Andersen, L. (2003). Identification of the C-signal, a contact-dependent morphogen coordinating multiple developmental responses in *Myxococcus xanthus*. Genes & development, 17(17), 2151–2161. doi:10.1101/gad.274203

Locascio, A., & Nieto, M. A. (2001). Cell movements during vertebrate development: integrated tissue behaviour versus individual cell migration. Current Opinion in Genetics & Development, 11(4), 464–469. doi:10.1016/S0959-437X(00)00218-5

MacArthur, B.D., Ma’ayan, A. & Lemischka, I.R. (2009). Systems biology of stem cell fate and cellular reprogramming. Nature Reviews Molecular cell biology, 10(10), 672–681. doi:10.1038/nrm2766

Maynard-Smith J. & Szathmáry E. (1995). The Major Transitions in Evolution. W.H. Freeman Spektrum.

Milligan, C. E., Prevette, D., Yaginuma, H., Homma, S., Cardwellt, C., Fritz, L. C., Tomaselli, K. J., Oppenheim, R. W. & Schwartz, L. M. (1995). Peptide inhibitors of the ICE protease family arrest programmed cell death of motoneurons *in vivo* and *in vitro*. Neuron, 15(2), 385–393. doi:10.1016/0896-6273(95)90042-X

Milo, R., Shen-Orr, S., Itzkovitz, S., Kashtan, N., Chklovskii, D. & Alon, U. (2002). Network motifs: simple building blocks of complex networks. Science, 298(5594), 824–827. doi:10.1126/science.298.5594.824

Monod, J. & Jacob, F. (1961) General conclusions: teleonomic mechanisms in cellular metabolism, growth, and differentiation. In Cold Spring Harbor symposia on quantitative biology, 26, 389–401. Cold Spring Harbor Laboratory Press.

Mouw, J. K., Ou, G., & Weaver, V. M. (2014). Extracellular matrix assembly: a multiscale deconstruction. Nature Reviews Molecular Cell Biology, 15(12), 771–785. doi:10.1038/nrm3902

Müller, F. D., Treuner-Lange, A., Heider, J., Huntley, S. M., & Higgs, P. I. (2010). Global transcriptome analysis of spore formation in *Myxococcus xanthus* reveals a locus necessary for cell differentiation. BMC Genomics, 11, 264. doi:10.1186/1471-2164-11-264

Müller, F. D., Schink, C. W., Hoiczyk, E., Cserti, E., & Higgs, P. I. (2012). Spore formation in *Myxococcus xanthus* is tied to cytoskeleton functions and polysaccharide spore coat deposition. Molecular Microbiology, 83(3), 486–505. doi:10.1111/j.1365-2958.2011.07944.x

Nanjundiah, V., Ruiz-Trillo, I., & Kirk, D. (2018). Protists and multiple routes to the evolution of multicellularity. Cells in Evolutionary Biology: Translating Genotypes into Phenotypes – Past, Present, Future, 71. doi:10.1201/9781315155968

Nariya, H., & Inouye, S. (2005). Identification of a protein Ser/Thr kinase cascade that regulates essential transcriptional activators in *Myxococcus xanthus* development. Molecular Microbiology, 58(2), 367–379. doi:10.1111/j.1365-2958.2005.04826.x

Nariya, H., & Inouye, S. (2006). A protein Ser/Thr kinase cascade negatively regulates the DNA-binding activity of MrpC, a smaller form of which may be necessary for the *Myxococcus xanthus* development. Molecular Microbiology, 60(5), 1205–1217. doi:10.1111/j.1365-2958.2006.05178.x

Nariya, H., & Inouye, M. (2008). MazF, an mRNA interferase, mediates programmed cell death during multicellular *Myxococcus* development. Cell, 132(1), 55–66. doi:10.1016/j.cell.2007.11.044

Newman, S. A., & Bhat, R. (2008). Dynamical patterning modules: physico-genetic determinants of morphological development and evolution. Physical Biology, 5(1), 015008. doi:10.1088/1478-3975/5/1/015008

Newman, S. A., & Bhat, R. (2009). Dynamical patterning modules: a “pattern language” for development and evolution of multicellular form. International Journal of Developmental Biology, 53(5-6), 693–705. doi: 10.1387/ijdb.072481sn

Ogawa, M., Fujitani, S., Mao, X., Inouye, S., & Komano, T. (1996). FruA, a putative transcription factor essential for the development of *Myxococcus xanthus*. Molecular Microbiology, 22(4), 757–767. 10.1046/j.1365-2958.1996.d01-1725.x

Perez-Carrasco, R., Barnes, C. P., Schaerli, Y., Isalan, M., Briscoe, J., & Page, K. M. (2018). Combining a Toggle Switch and a Repressilator within the AC-DC Circuit Generates Distinct Dynamical Behaviors. Cell Systems, 6(4), 521–530. doi:10.1016/j.cels.2018.02.008

Plamann, L., Kuspa, A., & Kaiser, D. (1992). Proteins that rescue A-signal-defective mutants of *Myxococcus xanthus*. Journal of Bacteriology, 174(10), 3311–3318. doi:10.1128/jb.174.10.3311-3318.1992

Pomerening, J.R., Sontag, E.D. & Ferrell Jr, J.E. (2003). Building a cell cycle oscillator: hysteresis and bistability in the activation of Cdc2. Nature Cell Biology, 5(4), 346–351. doi:10.1038/ncb954

Prill, R. J., Iglesias, P. A., & Levchenko, A. (2005). Dynamic properties of network motifs contribute to biological network organization. PLoS Biology, 3(11), e343. doi:10.1371/journal.pbio.0030343

Prochazka, L., Benenson, Y. & Zandstra, P.W. (2017). Synthetic gene circuits and cellular decision-making in human pluripotent stem cells. Current Opinion in Systems Biology, 5, 93–103. doi:10.1016/j.coisb.2017.09.003

Rivera-Yoshida, N., Del Angel, J. A. A., & Benitez, M. (2018). Microbial multicellular development: mechanical forces in action. Current Opinion in Genetics & Development, 51, 37–45. doi:10.1016/j.gde.2018.05.006

Rivera-Yoshida, N., Arzola, A. V., Arias Del Angel, J. A., Franci, A., Travisano, M., Escalante, A. E., & Benitez, M. (2019). Plastic multicellular development of *Myxococcus xanthus*: genotype–environment interactions in a physical gradient. Royal Society Open Science, 6(3), 181730. doi:10.1098/rsos.181730

Rolbetzki, A., Ammon, M., Jakovljevic, V., Konovalova, A., & Søgaard-Andersen, L. (2008). Regulated secretion of a protease activates intercellular signaling during fruiting body formation in *M. xanthus*. Developmental Cell, 15(4), 627–634. doi:10.1016/j.devcel.2008.08.002

Sager, B., & Kaiser, D. (1994). Intercellular C-signaling and the traveling waves of Myxococcus. Genes & Development, 8(23), 2793–2804. doi:10.1101/gad.8.23.2793

Shimkets, L. J., & Rafiee, H. (1990). CsgA, an extracellular protein essential for *Myxococcus xanthus* development. Journal of Bacteriology, 172(9), 5299–5306. doi:10.1128/jb.172.9.5299-5306.1990

Singer, M., & Kaiser, D. (1995). Ectopic production of guanosine penta-and tetraphosphate can initiate early developmental gene expression in *Myxococcus xanthus*. Genes & Development, 9(13), 1633–1644. doi:10.1101/gad.9.13.1633

Skotheim, J.M., Di Talia, S., Siggia, E.D. & Cross, F.R. (2008). Positive feedback of G1 cyclins ensures coherent cell cycle entry. Nature, 454(7202), 291–296. doi:10.1038/nature07118

Süel, G.M., Garcia-Ojalvo, J., Liberman, L.M. & Elowitz, M.B. (2006). An excitable gene regulatory circuit induces transient cellular differentiation. Nature, 440(7083), 545–550. doi:10.1038/nature04588

The Gene Ontology Consortium. (2019). The Gene Ontology Resource: 20 years and still GOing strong. Nucleic Acids Research, 47(D1), D330–D338. doi:10.1093/nar/gky1055.

Theocharis, A. D., Skandalis, S. S., Gialeli, C., & Karamanos, N. K. (2016). Extracellular matrix structure. Advanced Drug Delivery Reviews, 97, 4–27. doi:10.1016/j.addr.2015.11.001

Thompson, D.W. (1942). On growth and form. Cambridge University Press.

Thöny-Meyer, L., & Kaiser, D. (1993). *devRS*, an autoregulated and essential genetic locus for fruiting body development in *Myxococcus xanthus*. Journal of Bacteriology, 175(22), 7450–7462. doi:10.1128/jb.175.22.7450-7462.1993

Toda, S., Blauch, L.R., Tang, S.K., Morsut, L. & Lim, W.A. (2018). Programming self-organizing multicellular structures with synthetic cell-cell signaling. Science, 361(6398), 156–162. doi:10.1126/science.aat0271

Tomlin, C.J. & Axelrod, J.D. (2007). Biology by numbers: mathematical modelling in developmental biology. Nature Reviews Genetics, 8(5), 331–340. doi:10.1038/nrg2098

Tsai, T.Y.C., Choi, Y.S., Ma, W., Pomerening, J.R., Tang, C. & Ferrell, J.E. (2008). Robust, tunable biological oscillations from interlinked positive and negative feedback loops. Science, 321(5885), 126–129. doi:10.1126/science.1156951

Turing, A.M. (1952). The chemical basis of morphogenesis. Philosophical Transactions of the Royal Society of London Series B, 237, 37–72.

Udo, H., Munoz-Dorado, J., Inouye, M., & Inouye, S. (1995). *Myxococcus xanthus*, a gram-negative bacterium, contains a transmembrane protein serine/threonine kinase that blocks the secretion of beta-lactamase by phosphorylation. Genes & Development, 9(8), 972–983. doi:10.1101/gad.9.8.972

Ueki, T., & Inouye, S. (2003). Identification of an activator protein required for the induction of *fruA*, a gene essential for fruiting body development in *Myxococcus xanthus*. Proceedings of the National Academy of Sciences, 100(15), 8782–8787. doi:10.1073/pnas.1533026100

Verd, B., Monk, N. A., & Jaeger, J. (2019). Modularity, criticality, and evolvability of a developmental gene regulatory network. eLife, 8, e42832. doi:10.7554/eLife.42832.

Xiong, W. & Ferrell Jr, J.E. (2003). A positive-feedback-based bistable ‘memory module’ that governs a cell fate decision. Nature, 426(6965), 460–465. doi:10.1038/nature02089

Yang, Z. & Higgs, P. (2014). Myxobacteria: Genomics, Cellular and Molecular Biology. Caister Academic Press.

Zhang, W., Munoz-Dorado, J., Inouye, M., & Inouye, S. (1992). Identification of a putative eukaryotic-like protein kinase family in the developmental bacterium *Myxococcus xanthus*. Journal of Bacteriology, 174(16), 5450–5453. doi:10.1128/jb.174.16.5450-5453.1992

Zhu, J., Zhang, Y. T., Alber, M. S., & Newman, S. A. (2010). Bare bones pattern formation: a core regulatory network in varying geometries reproduces major features of vertebrate limb development and evolution. PLoS One, 5(5), e10892. doi:10.1371/journal.pone.0010892

